# Inactive variants of death receptor p75^NTR^ reduce Alzheimer’s neuropathology by interfering with APP internalization

**DOI:** 10.1101/2020.01.10.901926

**Authors:** Chenju Yi, Ket Yin Goh, Lik-Wei Wong, Kazuhiro Tanaka, Sreedharan Sajikumar, Carlos F. Ibáñez

## Abstract

A prevalent model of Alzheimer’s disease (AD) pathogenesis postulates the generation of neurotoxic fragments derived from the amyloid precursor protein (APP) after its internalization to endocytic compartments. However, the molecular pathways that regulate APP internalization and intracellular trafficking in neurons are unknown. Here we report that 5xFAD mice, an animal model of AD, expressing signaling-deficient variants of the p75 neurotrophin receptor (p75^NTR^) show greater neuroprotection from AD neuropathology than animals lacking this receptor. p75^NTR^ knock-in mice lacking the death domain or transmembrane Cys^259^ showed lower levels of Aβ species, amyloid plaque burden, gliosis, mitochondrial stress and neurite dystrophy than global knock-outs. Strikingly, long-term synaptic plasticity and memory, which are completely disrupted in 5xFAD mice, were fully recovered in the knock-in mice. Mechanistically, we found that p75^NTR^ interacts with APP and regulates its internalization in hippocampal neurons. Inactive p75^NTR^ variants internalized much slower and to lower levels than wild type p75^NTR^, favoring non-amyloidogenic APP cleavage by reducing APP internalization and colocalization with BACE1, the critical protease for generation of neurotoxic APP fragments. These results reveal a novel pathway that directly and specifically regulates APP internalization, amyloidogenic processing and disease progression, and suggest that inhibitors targeting the p75^NTR^ transmembrane domain may be an effective therapeutic strategy in AD.

## Introduction

A central tenet of the amyloid hypothesis of AD pathogenesis is the generation of neurotoxic fragments of APP through a series of proteolytic cleavages (Selkoe & Hardy, 2016; Karran *et al*, 2011). APP cleavage by BACE (beta-site APP cleaving enzyme) at an extracellular site close to the plasma membrane leaves a transmembrane C-terminal stub (beta-carboxyterminal fragment or CTFβ) that serves as a substrate for further intramembrane cleavage by gamma-secretase, a multisubunit complex that includes the aspartyl protease presenilin-1 (PS1) as its catalytic subunit. Cleavage by the gamma-secretase complex liberates a soluble CTFβ and a small N-terminal fragment of 40 or 42 amino acids in length known as the amyloid beta peptide or Aβ, the main component of the amyloid plaques that accumulate in the AD brain (Selkoe & Hardy, 2016; Karran *et al*, 2011). The majority of familial AD cases are caused by mutations in the genes encoding APP or PS-1 (Selkoe & Hardy, 2016; Karran *et al*, 2011), supporting the amyloid hypothesis of AD pathogenesis. APP can also be cleaved by cell surface alpha-secretases, most notably ADAM10, in an extracellular site very close to the plasma membrane, but C-terminal to the site of BACE cleavage. Thus, alpha-secretase cleavage precludes the generation of all BACE-derived products, including Aβ, and constitutes the non-amyloidogenic pathway in APP processing. Cleavage by alpha-secretase generates a soluble N-terminal fragment (sAPPα) and a C-terminal stub (sCTFα) that can be further processed by gamma-secretase.

Recent studies have indicated that, while cleavage by alpha secretases occurs at the plasma membrane, proteolytic processing by BACE requires APP internalization from the cell surface and thus mainly takes places in intracellular, endocytic compartments (Haass *et al*, 2012). Several studies have linked APP internalization to Aβ production (Koo & Squazzo, 1994; Selkoe *et al*, 1996), and complementary lines of evidence support this notion, including the requirement of low pH for BACE optimal catalytic activity (Vassar *et al*, 2009), the fact that genetic or pharmacological inhibition of APP internalization reduces Aβ generation (Carey *et al*, 2005), and evidence that neuronal activity enhances Aβ production by inducing APP internalization and trafficking to BACE-containing endosomes (Das *et al*, 2013; 2016). Thus, the localization and intracellular trafficking of APP appear to be critical for the balance between competing amyloidogenic and non-amyloidogenic pathways of APP processing. However, aside from classical components of the endocytic machinery, such as dynamin, our knowledge of the molecular pathways that can regulate APP internalization and intracellular trafficking in neurons is very rudimentary.

p75^NTR^ is a member of the death receptor superfamily, characterized by the presence of a death domain (DD) in their intracellular region (Liepinsh *et al*, 1997), which also includes the Tumor Necrosis Factor Receptor 1 (TNFR1), CD40, Fas and others (Ibáñez & Simi, 2012). Many of these receptors can induce cell death pathways as a mechanism for clearing damage produced after a lesion or insult. However, after severe injury or disease, they can also amplify tissue damage as a result of overactivation and/or overexpression. Upon neural injury or cellular stress, p75^NTR^ signaling can contribute to neuronal death, axonal degeneration and synaptic dysfunction (Ibáñez & Simi, 2012). p75^NTR^ can function as a receptor of neurotrophins, a family of neurotrophic growth factors, that includes nerve growth factor (NGF), brain-derived neurotrophic factor (BDNF) and others, as well as other ligands unrelated to the neurotrophins, including Aβ (for review, see (Underwood & Coulson, 2008)). Notably, expression of p75^NTR^ is increased in the brain of AD patients (Chakravarthy *et al*, 2012; Ernfors *et al*, 1990; Mufson & Kordower, 1992; Hu *et al*, 2002) as well as animal models of AD (Wang *et al*, 2011; Chakravarthy *et al*, 2010). Aβ can induce rapid cell death in cultured neurons through direct interaction with p75^NTR^ and downstream activation of cell death pathways (Perini *et al*, 2002; Sotthibundhu *et al*, 2008; Rabizadeh *et al*, 1994; Yaar *et al*, 1997; Knowles *et al*, 2009), but the relevance of *in vitro* overnight effects is unclear, as neuronal degeneration occurs during long periods of time in AD patients, and it is in fact seldom observed in animal models of AD. In line with this, elimination of p75^NTR^ affords only a mild improvement in those models (Knowles *et al*, 2009).

In this study, we used the 5xFAD model of AD (Oakley *et al*, 2006) to investigate neuropathological effects in different strains of knock-in and knock-out p75^NTR^ mutant mice. Unexpectedly, we found that knock-in mutations that inactivate p75^NTR^ signaling, but leave normal levels of receptor expression, conferred much higher protection from AD-associated neuropathology than a global knock-out. In our efforts to understand how an inactive receptor can afford greater neuroprotection than the absence of the receptor, we discovered a novel mechanism by which p75^NTR^ regulates APP internalization and its localization to intracellular compartments containing BACE.

## Results

### Reduced Aβ content and histopathology in the hippocampus of 5xFAD mice carrying inactive p75^NTR^ variants

The transmembrane domain of p75^NTR^ contains a highly conserved cysteine residue (Cys^259^ in mouse) which is critical for p75^NTR^ activation and signaling in response to neurotrophin ligands (Vilar *et al*, 2009). Neurons from knock-in mice carrying a Cys to Ala mutation at this position (C259A), or lacking the receptor death domain (ΔDD), are resistant to cell death induced by pro-neurotrophins and neurodegeneration induced by epileptic seizures, to a similar extent as knock-out (KO) neurons lacking p75^NTR^ entirely (Tanaka *et al*, 2016). Cortical neurons derived from these 3 lines also show comparable resistance to Aβ-mediated toxicity *in vitro* (Supplementary Figs. 1A-C). In order to study the contribution of p75^NTR^ activity to AD neuropathology *in vivo*, we crossed each of the 3 lines of p75^NTR^ mutant mice to the 5xFAD mouse model of AD. These mice express a human APP transgene carrying three mutations found in AD patients and a human PS-1 transgene with two AD mutations, both under regulatory sequences of a Thy1 transgene, thus primarily directing expression to neurons (Oakley *et al*, 2006). 5xFAD mice develop cerebral amyloid plaques starting at 2 months of age, achieve substantial Aβ burden with gliosis and neurite degeneration by 4 months, and show significant memory impairment by 6 months. The severity and accelerated progression of AD pathology displayed by this model served as a stringent test for assessing potential protective effects of the different p75^NTR^ mutations. At 5 months of age, 5xFAD mice showed increased levels of p75^NTR^ in the hippocampus compared to wild type mice, with prominent expression in dendrites of pyramidal neurons (Supplementary Fig. 1A). At 6 months, Western blots of hippocampal extracts indicated approximately 2-fold increase in p75^NTR^ protein levels in 5xFAD mice compared to wild type controls (Supplementary Fig. 1B).

5xFAD mice developed progressively increased Aβ plaque burden as detected histologically in sections through the hippocampus; at 12 months of age, 5% of the area of the hippocampus was occupied by Aβ plaques (Figs. 1A and B). No Aβ immunoreactivity could be detected at any age in wild type mice (not shown). In the 5xFAD background, all three p75^NTR^ mutants (5xFAD/KO, 5xFAD/C259A and 5xFAD/ΔDD, respectively) showed reduced levels of Aβ plaque burden at all the ages examined (6, 9 and 12 months) compared to 5xFAD mice expressing wild type p75^NTR^ (Fig. 1B). However, we noted that the two knock-in mutants afforded greater protection than the knock-out, with differences against 5xFAD/KO mice reaching statistical significance, specially at 9 and 12 months of age (Fig. 1B). Next, we quantified the levels of Aβ1-42 monomers, oligomers and fibrils by ELISA, after differential detergent and acid extraction from hippocampus of 9 month old mice. All three p75^NTR^ mutations significantly lowered the levels of Aβ1-42 species in 5xFAD mice; with, again, the knock-in variants showing a stronger effect than the knock-out, particularly the strain carrying the C259A allele (Fig. 1C).

**Figure 1.**
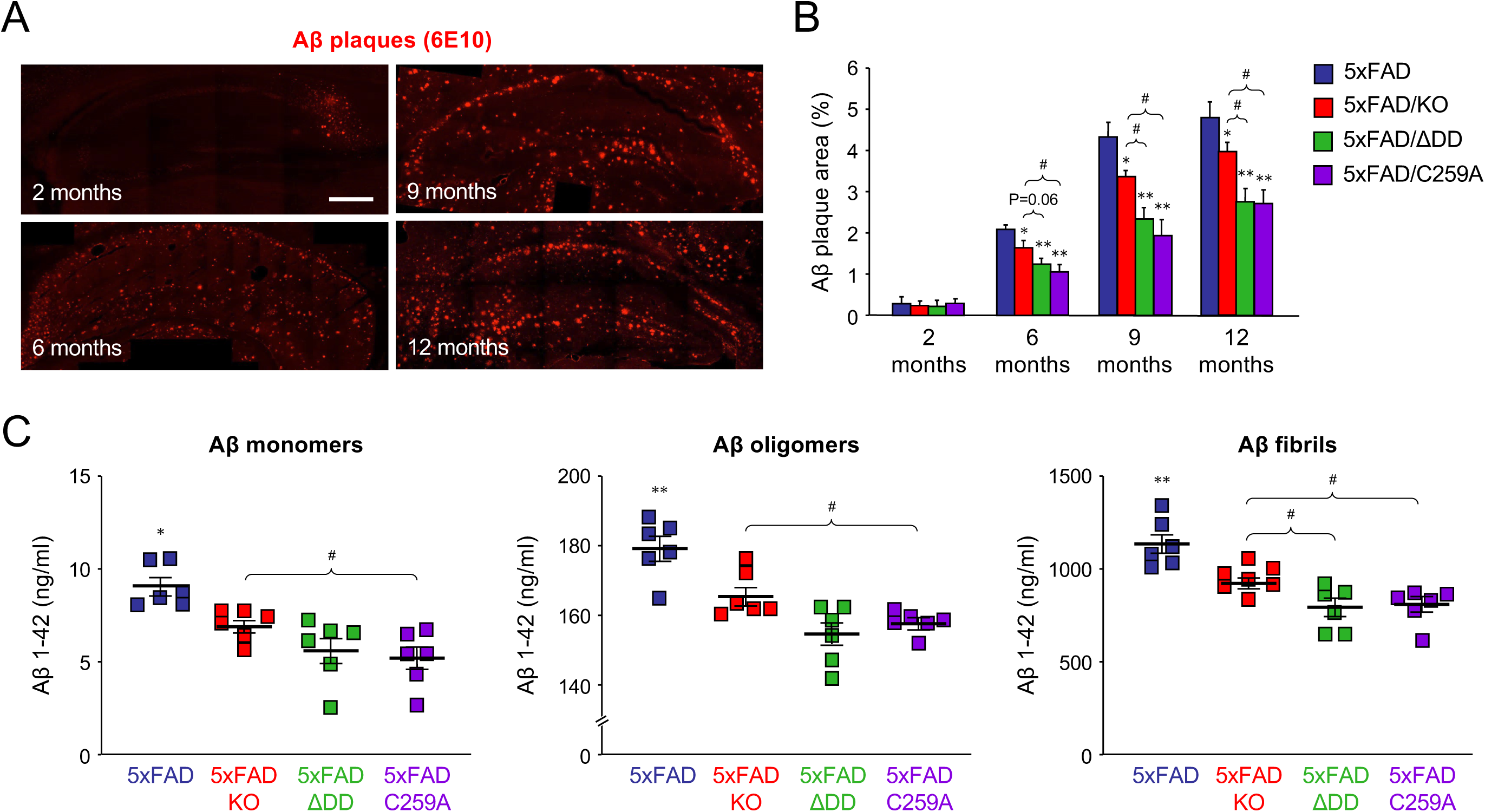
Reduced Aβ content in the hippocampus of 5xFAD mice carrying inactive p75^NTR^ variants. (A) Immunostaining for Aβ plaques with 6E10 antibody in coronal sections through the hippocampus of 5xFAD mice of the indicated ages. Scale bar, 400μm. (B) Quantification of Aβ plaque burden in the hippocampus of 5xFAD mouse strains carrying different p75^NTR^ variants as indicated. Histogram shows the percentage of hippocampal area occupied by Aβ plaques (average ± SEM, N=5 mice per group). *, P<0.05 and **, P<0.01 vs 5XFAD. #, P<0.05 vs 5XFAD/KO (one-way ANOVA followed by post hoc test). (C) ELISA determinations of Aβ1-42 content in hippocampus of 5xFAD mouse strains carrying different p75^NTR^ variants as indicated. Aβ monomers refers to the soluble fraction after TBS extraction, Aβ oligomers to the soluble fraction after RIPA buffer extraction of the TBS pellet, and Aβ fibrils to the soluble fraction after formic acid treatment of the RIPA pellet. See Materials and Methods for details.

Astrogliosis and microgliosis were significantly increased between 6 and 12 months of age in the hippocampus of 5xFAD mice; reaching 15-20% and 6-10% of the hippocampus area, respectively, compared to 5-10% and 2-4% in wild type controls (Figs. 2A-D). Astrocytes and microglial cells with morphology indicative of an activated state were found concentrated around Aβ plaques in the hippocampus of 5xFAD animals (Figs. 2A and C). All three p75^NTR^ mutations significantly reduced both forms of gliosis in the 5xFAD hippocampus. Again, the knock-in strains showed the lowest levels of gliosis, reaching statistically significant differences compared to 5xFAD/KO at 9 and 12 months, particularly in the 5xFAD/C259A strain (Figs. 2B and D). Next, we assessed the extent of neurite dystrophy in dendritic arbors of pyramidal hippocampal neurons, as assessed by accumulation of reticulon 3 (RTN3). Previous studies have shown RTN3 immunoreactivity to be markedly accumulated in dystrophic neurites in the brains of AD patients and APP transgenic mice (Hu *et al*, 2007). At 9 months, RTN3-positive neurites appeared as bright spots concentrated in and around Aβ plaques in the hippocampus of 5xFAD mice (Fig. 2E). No RTN3 immunoreactivity could be detected in wild type mice (not shown). RTN3 immunoreactivity in 5xFAD hippocampus was significantly reduced by all three p75^NTR^ mutations, with the strongest effects observed in the knock-in strains, particularly in 5xFAD/C259A mice (Fig. 2F). Finally, we looked at mitochondrial dysfunction, a well-known feature of AD neuropathology (Swerdlow, 2018), as assessed by MitoSOX staining, a mitochondrial superoxide indicator widely used to assess mitochondrial stress (Dikalov & Harrison, 2014). MitoSOX staining was significantly increased at 2 and 6 months in neurons of the pyramidal layer of the hippocampus of 5xFAD mice compared to wild type controls (Figs. 2G and H). At 2 months of age, all three p75^NTR^ mutations reduced MitoSOX levels significantly in the 5xFAD background, almost to the low levels found in wild type mice (Fig. 2H).

**Figure 2.**
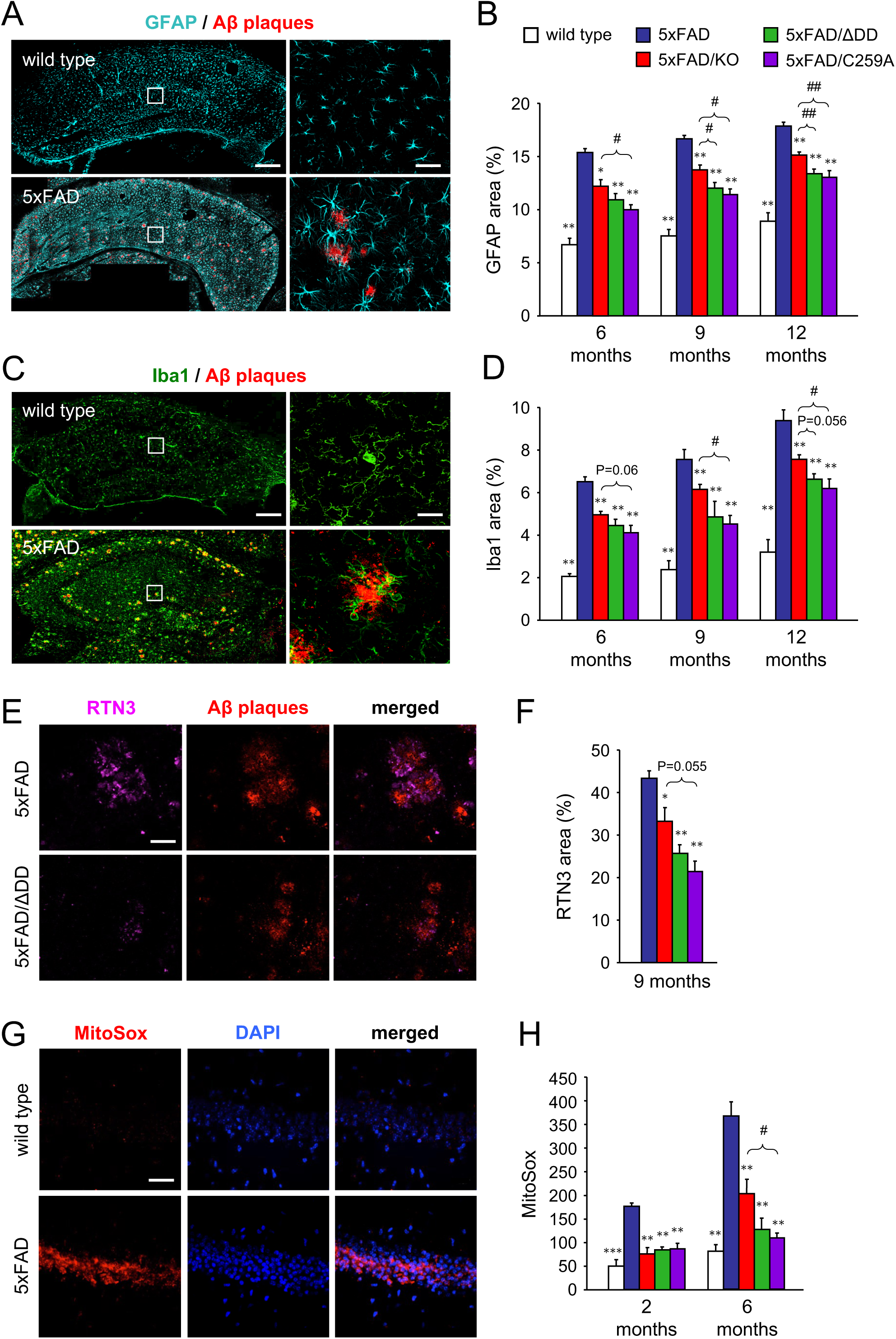
Reduced histopathology in the hippocampus of 5xFAD mice carrying inactive p75^NTR^ variants. (A) Immunostaining for Glial fibrillary acidic protein (GFAP), a marker of astrocytes, and Aβ plaques in coronal sections through the hippocampus of 6 month old wild type and 5xFAD mice. Scale bar, 300μm. Right hand panels show high magnification of the indicated areas. Scale bar, 50μm. (B) Quantification of GFAP area in the hippocampus of wild type and 5xFAD mouse strains carrying different p75^NTR^ variants as indicated. Histogram shows the percentage of hippocampal area occupied by GFAP immunostaining (average ± SEM, N=5 mice per group). *, P<0.05 and **, P<0.01 vs 5XFAD. #, P<0.05 and ##, P<0.01 vs 5XFAD/KO (one-way ANOVA followed by post hoc test). (C) Immunostaining for Ionized calcium binding adaptor molecule 1 (Iba1), a marker of microglial cells, and Aβ plaques in coronal sections through the hippocampus of 6 month old wild type and 5xFAD mice. Scale bar, 300μm. Right hand panels show high magnification of the indicated areas. Scale bar, 10μm. (D) Quantification of Iba1 area in the hippocampus of wild type and 5xFAD mouse strains carrying different p75^NTR^ variants as indicated. Histogram shows the percentage of hippocampal area occupied by Iba1 immunostaining (average ± SEM, N=5 mice per group). *, P<0.05 and **, P<0.01 vs 5XFAD. #, P<0.05 vs 5XFAD/KO. Other P values are indicated (one-way ANOVA followed by post hoc test). (E) Immunostaining of reticulon 3 (RTN3), a marker of dystrophic neurites, and Aβ plaques in coronal sections through the hippocampus of 6 month old 5xFAD and 5xFAD/ΔDD mice. Scale bar, 40μm. (F) Quantification of RTN3-positive dystrophic neurite area in the hippocampus of 5xFAD mouse strains carrying different p75^NTR^ variants as indicated. Histogram shows the percentage of Aβ plaque area that overlapped with RTN3 immunostaining (average ± SEM, N=5 mice per group). *, P<0.05 and **, P<0.01 vs 5XFAD. Other P values are indicated (one-way ANOVA followed by post hoc test). (G) MitoSox staining, a mitochondrial superoxide indicator, and DAPI in coronal sections through the hippocampus of 6 month old 5xFAD and 5xFAD/ΔDD mice. Scale bar, 60μm. (H) Quantification of MitoSOX signal in the pyramidal cell layer of hippocampus of wild type and 5xFAD mouse strains carrying different p75^NTR^ variants as indicated. Histogram shows MitoSox mean fluorescence intensity in arbitrary units (average ± SEM, N=5 mice per group). **, P<0.01 and ***, P<0.001 vs 5XFAD (one-way ANOVA followed by post hoc test).

However, at 6 months, there was a significant advantage of the p75^NTR^ knock-in strains over 5xFAD/KO mice, particularly in the 5xFAD/C259A strain, which showed MitoSOX levels indistinguishable from wild type and significantly lower than the knock-out (Fig. 2H).

Together, these studies indicated a significantly higher level of neuroprotection in mouse strains carrying signaling-deficient p75^NTR^ variants compared to knock-out mice lacking the receptor. This was in contrast to the results of the *in vitro* assay of Aβ neurotoxicity (Supplementary Figure 1), in which neurons from all three strains were equally resistant, suggesting that additional mechanisms must operate to account for the differences observed *in vivo*.

### Inactive p75^NTR^ variants, but not the knock-out, fully rescue synaptic deficits and memory impairment in 5xFAD mice

We then asked whether the beneficial effects on AD histopathology afforded by the different p75^NTR^ mutants had an impact on synaptic function and cognitive behavior of 5xFAD mice. Previous studies have shown that AD is associated with deficits in various forms of synaptic plasticity, including hippocampal long-term potentiation (LTP), in human patients and animal models (Mango *et al*, 2019; Selkoe, 2002; Palop & Mucke, 2010; Koch *et al*, 2012). In our studies, we used a form of late LTP induced by theta burst stimulation (TBS-LTP) in hippocampal area CA1. Hippocampal slices from wild type mice showed pronounced potentiation induced by TBS that was sustained for at least 4 hours (Fig. 3A). In contrast, synaptic potentiation was not maintained and declined rapidly in slices derived from 6 month old 5xFAD mice, indicating a profound deficit in LTP induction (Fig. 3A). Slices from 5xFAD/KO mice showed intermediate levels of potentiation that declined slowly over the course of the 4h recording, indicating a partial recovery (Fig. 3A). Remarkably, TBS-LTP in slices from 5xFAD/C259A and 5xFAD/ΔDD mice was strong in terms of induction, persistent, and maintained throughout the whole recording period of 4h, and was essentially indistinguishable from that recorded in wild type slices (Fig. 3A). Quantification of the change in field excitatory postsynaptic potential (fEPSP) revealed a stronger rescue of synaptic function in the p75^NTR^ knock-in strains, reaching statistically significant differences from 5xFAD mice throughout the recording period, unlike the knock-out (Fig. 3bB). We note that TBS-LTP in the absence of the 5xFAD transgene was normal in all three mutant strains and identical to that of wild type animals (Supplementary Figs. 3A and B).

**Figure 3.**
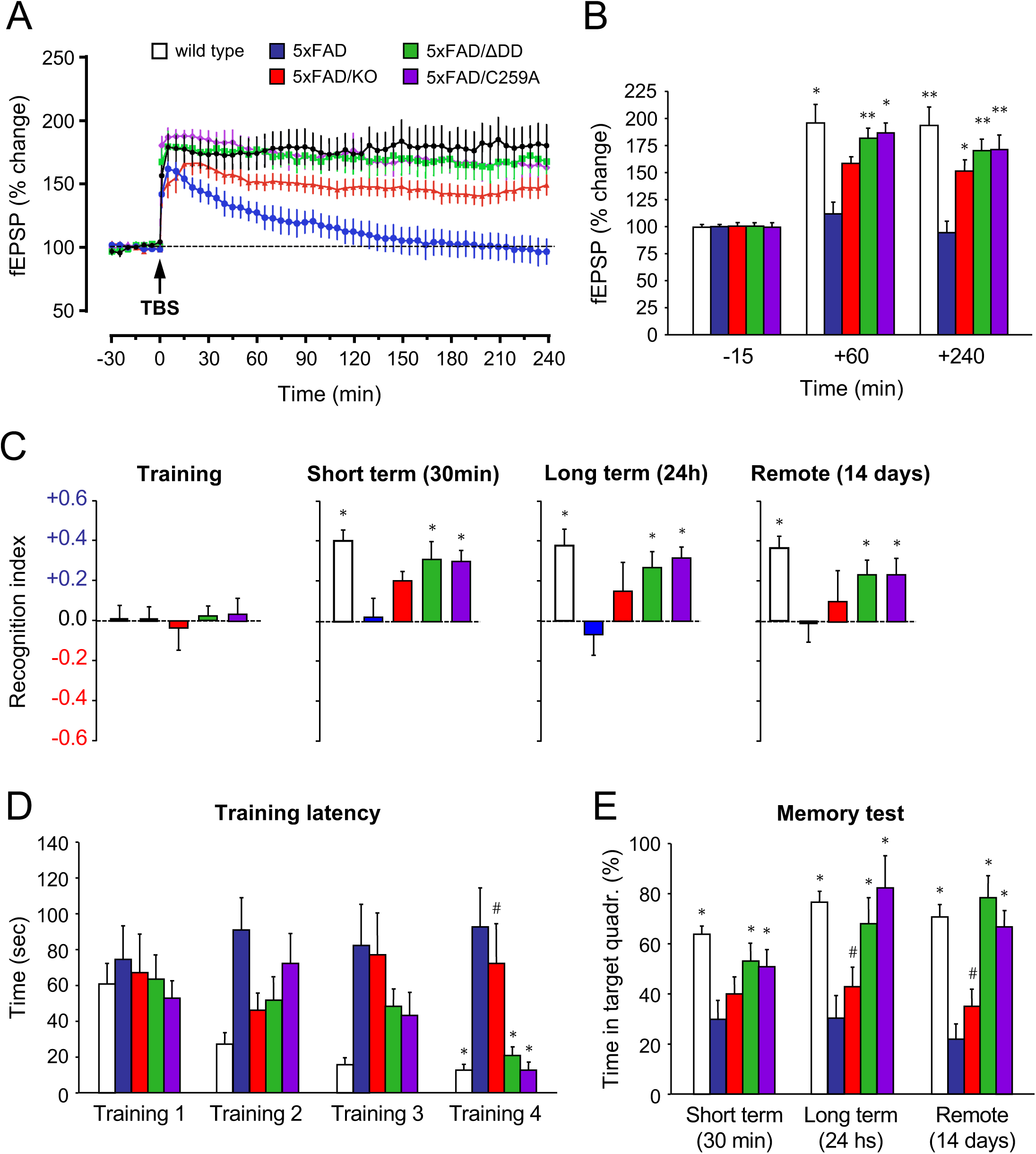
Inactive p75^NTR^ variants, but not the knock-out, fully rescue synaptic deficits and memory impairment of 5xFAD mice. (A) Percentage of change in field excitatory postsynaptic potential (fEPSP) recorded after theta-burst stimulation (TBS) in Schaffer collaterals of hippocampal slices from 6 month old wild type, 5xFAD and 5xFAD/p75^NTR^ mutant mice, as indicated. Results are presented as average % change normalized to t=0 ± SEM. N=6 (wild type), 5 (5xFAD), 9 (5xFAD/KO), 7 (5xFAD/ΔDD) and 8 (5xFAD/C259A) slices from 3 mice per genotype, respectively. (B) Quantification of fEPSP (average % change ± SEM) in the indicated genotypes at 3 time points. *, P<0.05; **, P<0.01 vs. 5xFAD (two-way ANOVA followed by post hoc test). N numbers as in (A). (C) Behavior in the novel object recognition (NOR) test of 6 month old wild type, 5XFAD and 5XFAD/p75^NTR^ mutant mice, as indicated. Histograms show average recognition index ± SEM during training, and 30min, 24h and 14 days after training, corresponding to measures of short term, long term and remote memory, respectively. Bar color codes are as in panel (A). *, P<0.05 vs. 5xFAD (two-way ANOVA followed by post hoc test). N=12 (wild type, 5xFAD/ΔDD and 5xFAD/C259A), 10 (5xFAD) and 8 (5xFAD/KO) mice per genotype, respectively. (D) Training latency in the Barnes maze test of 6 month old wild type, 5XFAD and 5XFAD/p75^NTR^ mutant mice, as indicated. Histograms show average latency in seconds to find the platform hole ± SEM in 4 consecutive training sessions. Bar color codes are as in panel (A). *, P<0.05 vs. 5xFAD; #, P<0.05 vs. wild type, 5xFAD/ΔDD or 5xFAD/C259A (two-way ANOVA followed by post hoc test). N=14 (wild type and 5xFAD/ΔDD), 10 (5xFAD, (5xFAD/KO and 5xFAD/C259A) mice per genotype, respectively. (E) Percentage of time spent in the target quadrant of the Barnes maze test 30min, 24h and 14 days after training. *, P<0.05 vs. 5xFAD; #, P<0.05 vs. wild type, 5xFAD/ΔDD or 5xFAD/C259A (two way ANOVA followed by post hoc test). N numbers as in (D).

Next, we assessed learning and memory in 6 month old animals using two different paradigms based on novelty and spatial memory, respectively, namely the novel object recognition test (NOR) and the Barnes maze. There was no difference between strains during training in the NOR test, when the animals are confronted with two identical objects, with recognition index (RI) close to zero, indicating no preference between the objects (Fig. 3C). Probe tests were conducted 30 min, 24h and 14 days after training, to assess short-term, long-term and remote memory, respectively, by confronting animals to a new object placed besides a familiar object. Wild type mice spent approximately 40% more time (RI≈0.4) on the new object at all three probe time points, which is the expected response of normal mice in the NOR test (Fig. 3C). In contrast, 5xFAD mice showed no indication that they recognized the new object (RI close to zero) at any time point (Fig. 3C), indicating severe memory disruption. Knock-out of p75^NTR^ afforded some degree of rescue, with average RI values between 0 and 0.15, but without reaching statistically significant differences compared to 5xFAD mice expressing p75^NTR^ at any time point, despite the large number of animals used (Fig. 3C). On the other hand, the deficits of 5xFAD mice were nearly completely rescued by either of the two p75^NTR^ knock-in alleles, showing average RI values that were not statistically different from those of wild type mice (Fig. 3C). The differences in cognitive performance between p75^NTR^ knock-in and knock-out strains were more striking the Barnes maze, a test of spatial memory strongly dependent on hippocampal function. During the training phase, wild type animals showed the expected learning performance, with progressively decreasing latencies in finding the escape hole, reaching 10 seconds or less by the 4^th^ training session (Fig. 3D). In contrast, 5xFAD mice showed no reduction of training latency, spending well over a minute to find the escape hole even after 4 training sessions (Fig. 3D), indicating a very strong learning deficit. As before, probe tests were conducted 30 min, 24h and 14 days after training to assess memory of the previous location of the escape hole. While wild type animals spent most (60-70%) of the probe time in the target quadrant, the performance of 5xFAD mice was not different from chance (≈25% in the target quadrant, Fig. 3E) at all time points, indicating no memory of the previous location of the escape hole. Remarkably, knock-out of p75^NTR^ did not confer any benefit to 5xFAD mice in this test. Latencies remained high and unchanged during training, and probe memory tests did not show statistically significant differences from 5xFAD mice expressing p75^NTR^ at any time point (Fig. 3E). In contrast, the p75^NTR^ knock-in alleles almost completely reverted the deficits of 5xFAD mice (Fig. 3E). These animals reached similar latencies as wild type mice during training, after perhaps a small delay, and spent most of the probe time (50-80%) in the target quadrant at all time points tested (Fig. 3E), indicating normal memory performance. We conclude from these studies that, although loss of p75^NTR^ afforded some level of protection from AD-associated histopathology and synaptic deficits —though not in cognitive performance, greater neuroprotection and, in fact, nearly full synaptic and behavioral recovery was only observed in 5xFAD mice that expressed the C259A and ΔDD variants.

### Signaling-deficient p75^NTR^ variants favor non-amyloidogenic processing of APP

The discrepancy between the behavior of signaling-deficient p75^NTR^ variants *in vitro* and *in vivo* prompted us to investigate additional mechanisms. We reasoned that the lower Aβ plaque burden and reduced levels of Aβ species in the brain of 5xFAD mice carrying p75^NTR^ variants could be due to reduced amyloidogenic APP processing in the mutants. In order to investigate this, we examined the levels of CTFβ, a product of APP cleavage by BACE, in 9 month old hippocampus of the different strains. This analysis revealed reduced levels of CTFβ in 5xFAD mice carrying mutant alleles of p75^NTR^ (Fig. 4A). Interestingly, 5xFAD/ΔDD and 5xFAD/C259A mice showed significantly lower levels of CTFβ than 5xFAD/KO mice, indicating reduced amyloidogenic APP cleavage in the knock-in strains. We note that expression of full-length APP was indistinguishable between all 5xFAD strains (Supplementary Fig. 4A), demonstrating comparable transgene expression levels. We assessed mRNA and protein levels of the beta-secretase BACE1 in hippocampal extracts of the four 5xFAD strains, but did not detect any significant differences (Supplementary Figs. 4B and C). We reasoned that reduced amyloidogenic cleavage could have been due to increased non-amyloidogenic processing by alpha-secretase. We therefore assessed the levels of sAPPα, a product of the competing, non-amyloidogenic pathway. We found significantly increased levels of sAPPα in hippocampal extracts of 5xFAD mice carrying mutant p75^NTR^ alleles compared to 5xFAD animals expressing wild type p75^NTR^ (Fig. 4B). Mirroring the effects observed on CTFβ, 5xFAD/ΔDD and 5xFAD/C259A mice showed significantly higher levels of sAPPα than 5xFAD/KO mice, indicating that non-amyloidogenic APP cleavage is more prevalent in the knock-in strains. mRNA and protein levels of ADAM10, the main alpha-secretase, in hippocampal extracts of the four 5xFAD strains were comparable (Supplementary Figs. 4D and E). Together, these results indicated a bias favoring non-amyloidogenic alpha-processing of APP in 5xFAD strains carrying signaling-deficient alleles of p75^NTR^, without altered levels of the main secretases or the substrate.

**Figure 4.**
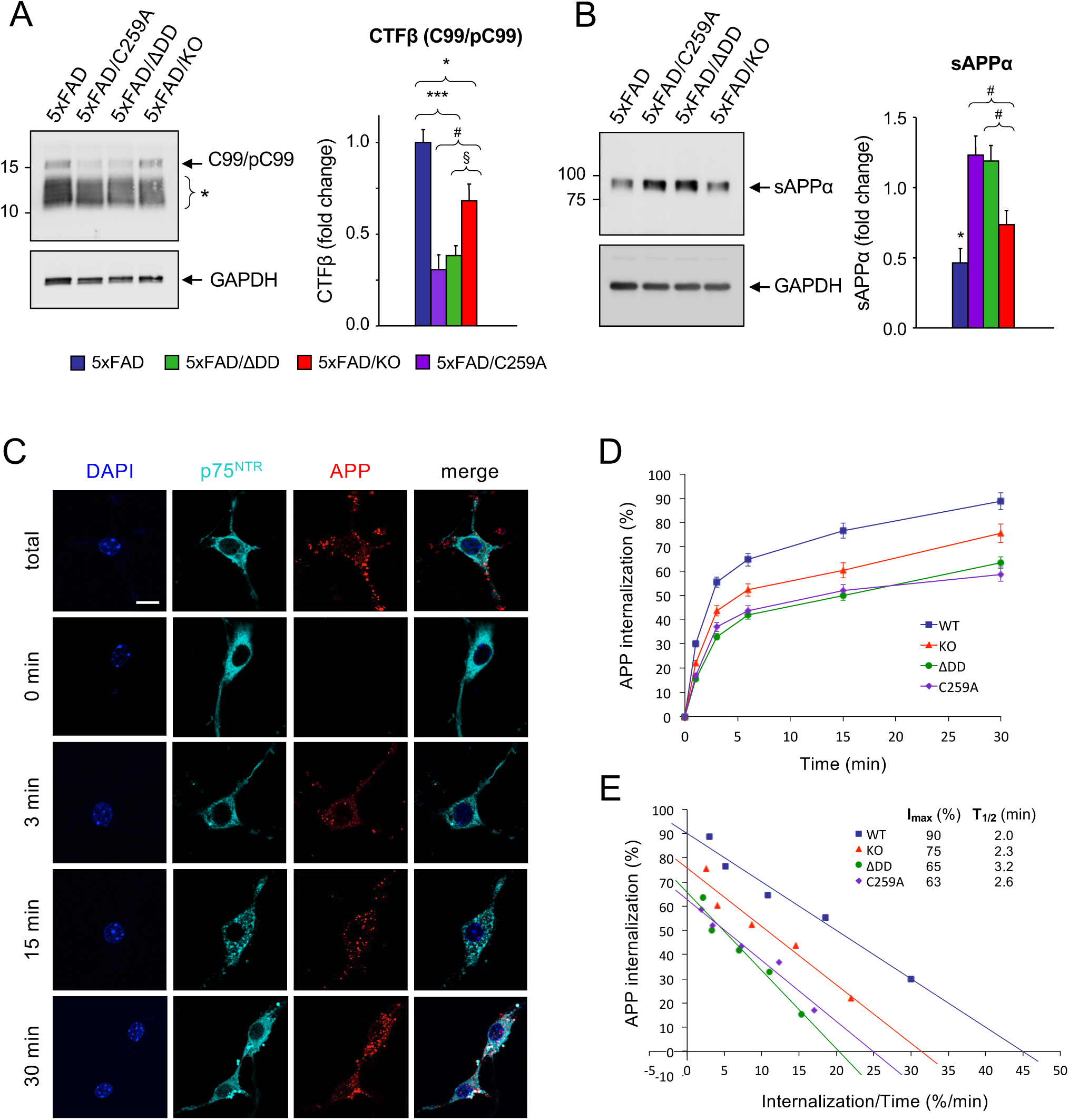
Reduced amyloidogenic processing and APP internalization in hippocampal neurons carrying signaling-deficient p75^NTR^ variants. (A) Western blot analysis of CTF beta (CTFβ) in hippocampal lysates of 9 month old 5xFAD mice carrying different p75^NTR^ alleles detected using anti-APP-CTF antibody (A8717, Table S1). Arrow points to C99/pC99 CTFβ species as previously assigned by (Buxbaum *et al*, 1998) and (Kwart *et al*, 2019). Bracket (*) denotes different species of native and phosphorylated alpha and beta CTFs of lower molecular weights (based on Figs. 3F and 5A in (Kwart *et al*, 2019)). Lower panel shows reprobing for GAPDH. Quantification to the right shows average ± SEM of C99/pC99 CTFβ species, normalized to GAPDH and expressed relative to levels in 5xFAD mice. *, P<0.05 vs, 5xFAD/KO; ***, P<0.001 vs. 5xFAD/ΔDD and 5xFAD/C259A; #, P<0.05 vs. 5xFAD/KO; §, P=0.056 vs. 5xFAD/KO. N=5 mice per group. (B) Western blot analysis of soluble APP alpha (sAPPα) in hippocampal lysates of 9 moth old 5xFAD mice carrying different p75^NTR^ alleles detected using anti APP 6E10 antibody (Table S1). Lower panel shows reprobing for GAPDH. Quantification to the right shows average ± SEM normalized to GAPDH and expressed relative to levels in 5xFAD mice. *, P<0.05 vs. 5xFAD/KO, 5xFAD/ΔDD and 5xFAD/C259A; #, P<0.05 vs. 5xFAD/KO. N=9 mice per group. (C) Internalization of 5xFAD hAPP in wild type mouse hippocampal neurons. Live neuron cultures were fed with anti-human APP antibodies (6E10) on ice, washed, then placed at 37°C for different periods of time to allow internalization. The reaction was stopped by a quick acid wash followed by fixation. Total staining (100%) was determined by direct fixation after antibody feeding. Baseline (t=0 min) was obtained by acid wash directly after antibody feeding. Counterstaining for p75^NTR^ (antibody GT15057) and DAPI are also shown. Scale bar, 10μm. (D) Internalization of hAPP in hippocampal neurons from wild type, p75^NTR^ knock-out, ΔDD and C259A mice. Show are averages ± SEM of percentage internalization of total surface APP (set to 100%). N=3 independent experiments each performed in duplicate. (E) Linear transformation of hAPP internalization kinetics shown in (D).

### Reduced APP internalization in hippocampal neurons carrying inactive p75^NTR^ variants

Realizing that the competing alpha- and beta-cleavage pathways of APP processing are known to occur in different subcellular compartments, namely plasma membrane versus intracellular endosomes, respectively, we hypothesized that perhaps substrate availability, rather than overall substrate or secretase levels, may underlie the bias favoring non-amyloidogenic APP processing in p75^NTR^ mutant mice. Following this line of thought, we reasoned that changes in APP internalization, and thus its steady-state residence at the plasma membrane, could differentially affect the access of alpha- and beta-secretases to APP. In order to test this idea, we established cultures of hippocampal neurons infected with a lentivirus expressing human APP carrying the 3 mutations found in 5xFAD mice. This afforded us a higher throughput of culture preparations compared to isolating neurons directly from 5xFAD embryos, as well as allowing control of APP specificity in PLA experiments (see below). Live neuron cultures were fed with anti-human APP antibodies on ice, then placed at 37°C for different periods of time to allow internalization (Fig. 4C; see Materials and Methods for details). In wild type neurons, maximal internalization of cell surface APP was close to 100% (I_max_≈90%) and relatively fast (T_1/2_≈2 min; Figs. 4D and E). In p75^NTR^ knock-out neurons, overall APP internalization was lower (I_max_≈75%) but of comparable speed (T_1/2_≈2.3 min; Figs. 4D and E). Interestingly, maximal APP internalization was further reduced in hippocampal neurons derived from ΔDD and C259A knock-in mice (I_max_≈65% and 63%, respectively) and considerably slower than in wild type neurons (T_1/2_≈3.2 and 2.6 min, respectively; Fig. 4E). These results suggested that p75^NTR^ is a positive regulator of APP internalization. The fact that APP internalization was even lower in neurons expressing signaling-deficient variants of p75^NTR^ suggested that receptor activity contributes to the effects of p75^NTR^ on APP internalization.

Noting that APP internalization was less efficient in ΔDD and C259A neurons than in KO neurons, we considered the possibility that, although signaling-deficient, the ΔDD and C259A variants may affect APP internalization by other means. To begin addressing this, we first evaluated the internalization of p75^NTR^ itself in hippocampal neurons derived from wild type, ΔDD and C259A mice, respectively, applying similar methodology as above, but using anti-mouse p75^NTR^ antibodies instead (Supplementary Fig. 5A). Maximal internalization of cell surface p75^NTR^ was close to saturation for wild type as well as ΔDD and C259A variants (I_max_≈106%, 96% and 94%, respectively; Supplementary Figs. 5B and C). In contrast, ΔDD and C259A were internalized at much slower speeds, approximately half, compared to wild type p75^NTR^ (T_1/2_≈12.8 and 11.1 min, respectively, compared to 5.4 min in wild type neurons; Supplementary Fig. 5C).

### APP and p75^NTR^ interact and co-internalize in hippocampal neurons

The slower internalization speeds of ΔDD and C259A p75^NTR^ correlated with the overall lower levels of APP internalization observed in neurons expressing these variants, prompting us to consider whether ΔDD and C259A may interfere with APP internalization by holding up APP molecules through direct interaction. An earlier study had reported that APP and p75^NTR^ can be co-immunoprecipitated from transfected cells and total brain extracts (Fombonne *et al*, 2009), but left open the question of whether they can interact under more physiological conditions on the plasma membrane of living neurons. To test this idea, we assessed whether APP and p75^NTR^ could interact directly on the surface of hippocampal neurons using the proximity ligation assay (PLA) (Supplementary Fig. 6A). We found identical levels of APP/p75^NTR^ complexes on the surface of neurons expressing either wild type, ΔDD or C259A p75^NTR^ (Supplementary Fig. 6B), indicating that all three receptor variants are able to interact with APP to the same extent. We did not detect any PLA signals in wild type neurons that were not infected with hAPP-expressing lentivirus, nor p75^NTR^ knock-out neurons expressing hAPP (Supplementary Figs. 6A and B).

Having demonstrated that APP and p75^NTR^ can interact directly in hippocampal neurons, we sought to determine whether the two molecules can also internalize together. To this end, we adapted the internalization assay to monitor trafficking of PLA signals arising from surface complexes co-labeled with anti-hAPP and anti-mouse p75^NTR^ antibodies (Fig. 5A). The internalization kinetics of complexes between APP and wild type p75^NTR^ (as detected by PLA) was very similar to that of total APP, with I_max_≈86% and T_1/2_≈2.2 min (Figs. 5B and C). In contrast, internalization of APP/p75^NTR^ complexes was markedly reduced, as well as slower, in neurons derived from ΔDD and C259A mice, with I_max_≈68 and 66% and T_1/2_≈4.3 and 3.9 min, respectively (Fig. 5C).

**Figure 5.**
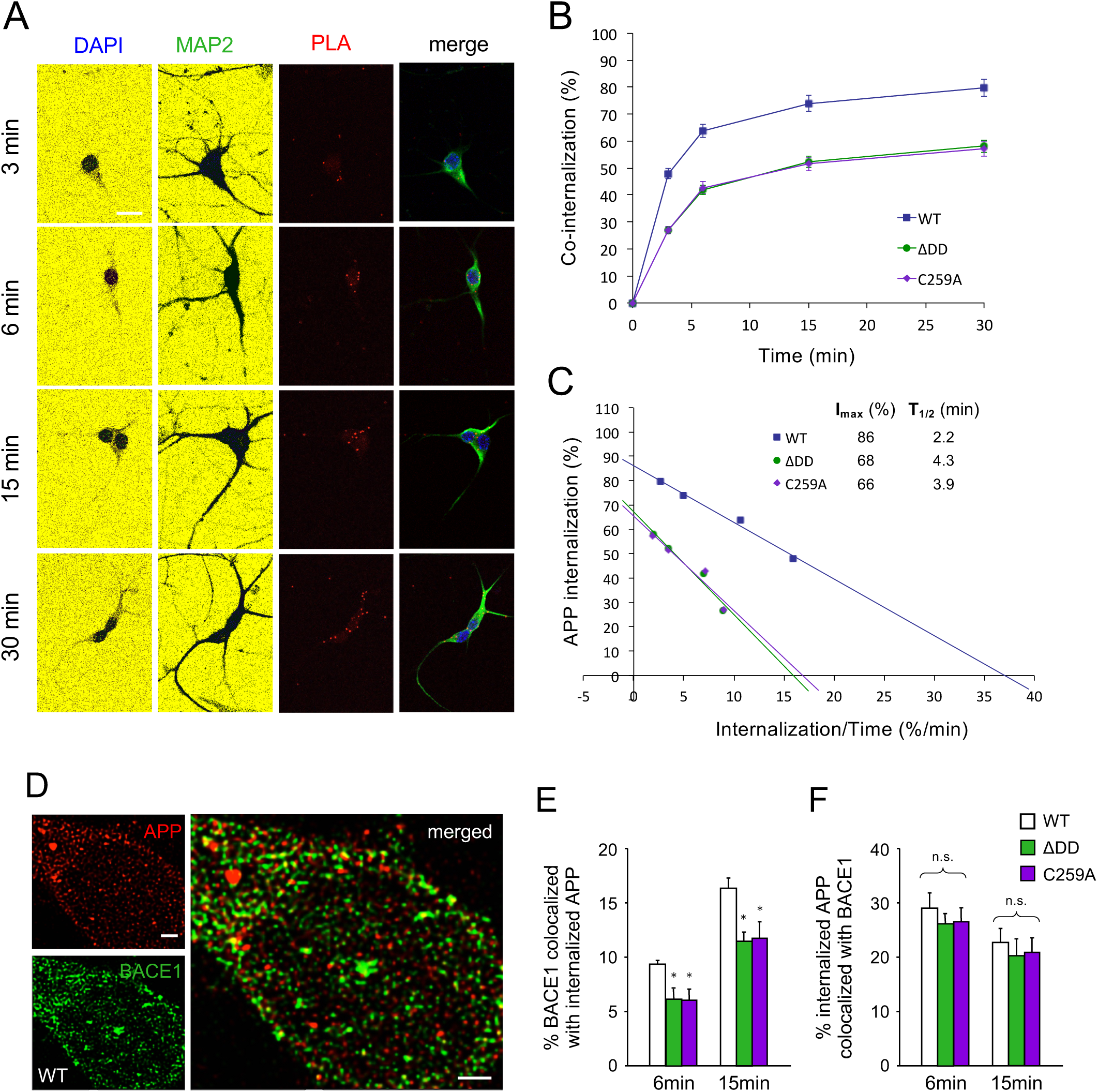
Reduced APP/p75^NTR^ co-internalization and trafficking of internalized APP to intracellular compartments containing BACE in neurons expressing ΔDD and C259A alleles. (A) Internalization of hAPP/p75^NTR^ PLA signals in wild type mouse hippocampal neurons. Live neuron cultures were fed with anti-mouse p75^NTR^ and anti-hAPP antibodies on ice, washed, then PLA reaction was performed and plates placed at 37°C for different periods of time to allow internalization of PLA signals. Counterstaining for MAP2 and DAPI are also shown. Scale bar, 10μm. (B) Co-internalization of hAPP and p75^NTR^ in hippocampal neurons from wild type, ΔDD and C259A mice. Show are averages ± SEM of percentage internalization of total surface PLA signal (set to 100%). N=3 independent experiments each performed in duplicate. (C) Linear transformation of p75^NTR^ internalization kinetics shown in (B). (D) Super-resolution micrographs of hAPP after 15min internalization (red), BACE1 immunocytochemistry (green) and their superimposition (merged) in wild type hippocampal neurons infected with 5xFAD hAPP lentivirus. Scale bar, 1μm. (E) Quantification of the proportion of BACE1 that co-localized with hAPP after 6 and 15 min of internalization at 37°C. Results are expressed as average ± SEM of % BACE co-localized with hAPP. N=3 independent experiments, each performed in duplicate; *, P<0.05 vs. WT. (F) Quantification of the proportion of internalized hAPP that co-localized with BACE1 after 6 and 15 min of internalization at 37°C. Results are expressed as average ± SEM of % internalized hAPP co-localized with BACE. N=3 independent experiments, each performed in duplicate; n.s., not significantly different.

### Reduced APP trafficking to intracellular compartments containing BACE1 in neurons expressing ΔDD and C259A p75^NTR^ alleles

Finally, we investigated whether decreased APP internalization in neurons expressing mutant ΔDD and C259A p75^NTR^ alleles translated into reduced encounters between APP and BACE in intracellular compartments, the first step in the amyloidogenic pathway. To this end, we used super-resolution microscopy to assess the extent to which internalized hAPP was localized to endosomes containing BACE1 in hippocampal neurons expressing different p75^NTR^ variants. Internalized hAPP was labeled as before, by feeding live neuron cultures infected with 5xFAD hAPP lentivirus with anti-hAPP antibodies, then allowing this label to traffic intracellularly at 37°C. This was followed by immunostaining for BACE1 (Fig. 5D). We found significantly lower levels of BACE1 co-localized with internalized hAPP in ΔDD and C259A p75^NTR^ mutant neurons compared to wild type neurons (Fig. 5E). When normalized to the levels of internalized APP, which are lower in the mutant neurons as indicated earlier, hAPP/BACE1 co-localization was not significantly different between genotypes (Fig. 5F), indicating that, once internalized, a comparable fraction of hAPP ends up in BACE-containing endosomes of wild type, ΔDD and C259A p75^NTR^ neurons.

## Discussion

In this study, we describe a new pathway that directly and specifically regulates APP internalization, its amyloidogenic processing and its impact on disease progression. Death receptor p75^NTR^ contributes to AD neuropathology by promoting internalization of APP to intracellular compartments containing BACE. p75^NTR^ interacts directly with APP at the cell surface, and enables higher levels of APP internalization compared to neurons lacking the receptor. A striking finding of this study, however, is the ability of signaling-deficient variants of p75^NTR^ to confer greater neuroprotection than the elimination of the receptor. This unexpected result can be explained by the significantly reduced internalization of these variants, which, through their interaction with APP, leads to reduced APP internalization, enhanced APP cleavage by alpha secretases, decreased APP trafficking to BACE1-containing endosomes, and decreased cleavage by beta secretase. We note that ΔDD and C259A could also reduce Aβ plaque burden by other mechanisms, including clearance of Aβ peptides (Wang *et al*, 2011), but such mechanism would also be present in neurons expressing wild type p75^NTR^, and hence less likely to account for the differences observed here. Moreover, the reduced levels of CTFβ and increased levels of sAPPα observed in 5xFAD/ΔDD and 5xFAD/C259A mice suggest reduced amyloidogenic processing, rather than enhanced clearance, as the predominant mechanism of Aβ plaque reduction in these mutants.

Several reports have linked the intracellular accumulation of CTFβ to early neurodegenerative processes in AD (Xu *et al*, 2016; Lauritzen *et al*, 2016; Pera *et al*, 2017; Lauritzen *et al*, 2012). We found that the ΔDD and C259A alleles of p75^NTR^ significantly reduced the accumulation of this fragment in the hippocampus of 5xFAD mice, which, together with decreased Aβ plaque formation, could also contribute to their neuroprotective effects. Interestingly, the levels of CTFβ and Aβ in the hippocampus of 5xFAD mice carrying different p75^NTR^ alleles also correlated with the extent of APP internalization. This novel mechanism is independent of the ability of p75^NTR^ to mediate some of the neurotoxic effects of Aβ oligomers, including cell death and neurite degeneration. However, these *in vitro* effects fail to explain both the time course as well as several aspects of the disease observed in AD patients and animal models.

Neurotrophin binding enhances p75^NTR^ internalization (Bronfman *et al*, 2003), underlying the important role of receptor signaling in intracellular trafficking. Hippocampal neurons produce neurotrophins endogenously, particularly BDNF, so the reduced internalization of ΔDD and C259A p75^NTR^ variants is likely linked to their inability to signal in response to endogenous ligands. Taken to its logical conclusion, such notion has an interesting corollary. Through their effects on p75^NTR^ internalization, neurotrophins may enhance APP intracellular trafficking to BACE endosomes, and hence Aβ production. It is also interesting to note that BDNF is well known for its ability to enhance neuronal activity (Lu *et al*, 2013), which, in turn, as shown by several recent studies, increases Aβ generation (Das *et al*, 2013; 2016).

The 5xFAD mouse model of AD displays enhanced and accelerated AD-like neuropathology and is perhaps one of the most aggressive AD models in mice. Importantly, however, 5xFAD mice do not display any abnormality that is not found in the AD patient population. Remarkably, we found a considerable reduction in the histopathology and nearly complete recovery of synaptic function and cognitive behavior in this rather strong model of AD after deletion of the p75^NTR^ death domain or mutation of its transmembrane Cys^259^. We find quite striking that changing a single amino acid in the mouse genome can have such dramatic effects on the course of this disease. The majority of previous efforts to identify small molecules targeting p75^NTR^ have focused on the extracellular domain of the receptor, with the intent of either mimicking or inhibiting neurotrophin binding. This has proven difficult, due to the large interfaces involved. The results of the present study suggest that targeting the transmembrane domain of p75^NTR^ may be a more promising strategy. We have recently provided proof-of-principle of this general concept in a recent report (Goh *et al*, 2018), paving the way for larger scale screenings of compound collections that may enable the discovery of substances mimicking the effects of the C259A mutation.

In summary, the results of the present study reveal an unexpected mechanism by which p75^NTR^ affects the generation of neurotoxic APP fragments through its ability to interact with APP and regulate APP internalization to intracellular compartments containing the amyloidogenic protease BACE. The fact that a single point mutation in a gene other than APP can have such strong neuroprotective effects in an aggressive model of AD highlights the importance of the transmembrane domain in the activation and function of p75^NTR^, and should encourage efforts to target this mechanism as a means to limit neurodegeneration in AD.

## Materials and Methods

### Animals

Mice were housed in a 12 hour light-dark cycle, and fed a standard chow diet. The mouse lines utilized in this study have been described previously and are as follows: 5xFAD (Oakley *et al*, 2006); p75^NTR^ exon 3 knock-out (Lee *et al*, 1992); and ΔDD and C259A p75^NTR^ knock-in mice with deletion of the death domain or a Cys^259^Ala substitution, respectively (Tanaka *et al*, 2016). All strains were back-crossed for at least 10 generations to a C57BL/6J background (considered as wild type). All animal procedures were approved by the National University of Singapore Institutional Animal Care and Use Committee.

### Primary culture of cortical and hippocampal neurons

Pregnant female mice were euthanized on the 17th day of gestation by injection of sodium pentobarbital followed by cervical dislocation. Cerebral cortical or hippocampal structures were aseptically removed from the embryos and digested in Papain (Sigma Aldrich) for 30 min at 37°C and rinsed in neuronal maintenance media. Neurons were triturated into a single cell suspension, counted with a hemocytometer, then transferred to coverslips coated with 0.01% poly-D-lysine (Sigma Aldrich) and 1μg/mL mouse Cultrex® Laminin (R&D Systems). Cultured hippocampal neurons were maintained in serum-free defined Neurobasal media supplemented with B27 (Invitrogen), GlutaMAX (Invitrogen) and 50 μg/mL Gentamicin (Invitrogen) at 37°C in 5% CO2.

### Lentivirus generation and transduction

HEK293FT cells (Invitrogen) were maintained in OptiMEM with 5% fetal bovine serum and antibiotics (Invitrogen). Cells were co-transfected with lentiviral constructs for overexpression with the packaging vectors Δ8.9 and VSV-G (Addgene) using FuGENE®-6 (Promega). Supernatants containing viral particles were aseptically collected 3 days post-transfection, filtered using 0.4μm PES filters (Sartorius), concentrated 50-100 times and dialyzed into sterile Dulbecco’s phosphate buffer saline (DPBS) by centrifugation in 100kDA cut-off Amicon Ultra-15 centrifugal filters (Millipore). Viruses were aliquoted and snap-frozen in liquid-nitrogen. Viruses were titrated by serial dilution on primary dissociated cortical neurons and quantified for EGFP expression with a Ti-E inverted fluorescence microscope. Lentiviral transduction of hippocampal neurons was performed after 1 day in vitro (DIV1) at a multiplicity of infection (MOI) =5 and left for 24h before the media was changed. The infected cultures were used at 5 days post-infection.

### Protein fractionation from mouse hippocampus and Aβ ELISA

For extraction of Aβ monomers, oligomers and fibrils, we followed a fractionation protocol previously described (Sherman & Lesné, 2011) with modifications as follows. Dissected frozen mouse hippocampal tissue (9-months old) was weighed, thawed on ice and homogenized using a manual Dounce homogenizer, in Tris-buffered saline (TBS, 25mM Tris, 140mM NaCl, pH 7.2-7.6) containing protease and phosphatase inhibitor (Nacalai Tesque) at a ratio of 1:9 (tissue:buffer, w/v).

Homogenates were then ultracentrifuged (Himac CS150GXL, Hitachi) at 100,000g for 1h at 4°C. The supernatant, which constitutes the soluble materials containing Aβ monomers, was collected as the TBS fraction. The pellet of the TBS fraction was resuspended in RIPA buffer (50mM Tris, pH 8, 150mM NaCl, 1% NP40, 5mM EDTA, 0.5% sodium deoxycholate, 0.1% SDS) containing protease and phosphatase inhibitors by trituration 10-15 times with a 1ml pipette followed by a 27G syringe needle. Samples were incubated at 4°C for 1h with agitation, and ultracentrifuged at 100,000g for another hour at 4°C. The supernatant constituted the RIPA fraction containing Aβ oligomers. The pellet of the RIPA fraction was dissolved in 70% formic acid (FA) and the homogenate was sonicated on ice for 30s (VCX 130, Sonics & Materials Inc; amplitude 40%, 3 x 10s of sonication). Sonicated samples were subjected to ultracentrifugation at 100,000g for 30 minutes at 4°C. The supernatant was collected and brought to neutral pH using neutralization buffer (1M Tris, 0.5M Na2HPO4, 0.05% NaN3) at 1:20 dilution factor. These samples constituted the FA fraction containing Aβ fibrils and were stored at room temperature to avoid precipitation at lower temperature. The TBS and RIPA fractions were stored at −20°C until further analysis. The content of Aβ1-42 in the TBS, RIPA and FA fractions was assessed using the Human Aβ1-42 enzyme-linked immunosorbent assay (ELISA) Kit (Merck) according to the manufacturer’s instructions. The concentration of Aβ1-42 was determined by the absorbance value detected using a microplate reader (BioTek Cytation 5, US) at 450 nm and 590 nm.

### Western blotting

Tissue lysates and protein fractions were mixed with 5X sample buffer (250mM Tris-HCl pH6.8, 10% SDS, 30% glycerol, 5% β-mercaptoethanol, 0.02% bromophenol blue) and boiled at 95°C for 5 min before electrophoresis on polyacrylamide gels. 16.5% Tris-Tricine gels (Biorad) were used to resolve APP CTF fragments. Proteins were blotted on polyvinylidene fluoride (PVDF) membranes (0.2µm; Amersham). Membranes were blocked using 5% non-fat milk (Biorad) in TBST (0.1% Tween-20) and incubated overnight at 4°C with primary antibodies as listed in Supplementary Table S1. Immunoreactivity was visualized using horseradish peroxidase (HRP)-conjugated secondary antibodies at 1:10,000 dilution. Immunoblots were developed using the SignalFire ECL Reagent (Cell Signaling) or SuperSignal West Femto (Pierce) and exposed to CL-Xposure Film (Thermo Fisher). Densitometric analysis of x-ray films was done using ImageJ software (NIH).

### Immunocytochemistry and immunohistochemistry

Hippocampal neurons cultured on coverslips were briefly washed in PBS, fixed for 15min in 4% paraformaldehyde solution (Sigma Aldrich) and blocked in PBS containing 0.2% gelatin and 0.25% Triton-X-100. Fixed cells were then incubated overnight at 4°C with the appropriate antibodies as listed in Supplementary Table S1, followed by incubation in fluorophore-conjugated secondary antibodies. Coverslips were mounted onto microscopy slides using fluoromount-G (SouthernBiotech). Confocal laser scanning microscopy was performed on a Leica SP8 microscope. For immunohistochemistry, cryostat sections (30 μm) of mouse brain prepared as previously described were fixed 30 min at room temperature with 4% paraformaldehyde (PFA), washed in PBS, treated 10 min with 70% formic acid (when mouse 6E10 was used) and preincubated in PBS containing 0.2% gelatin and 1% Triton-X-100 for 30 min. Sections were then processed for immunostaining by overnight incubation at 4 °C with primary antibodies as indicated in in Supplementary Table S1, washed in PBS and incubated in secondary donkey anti-rabbit and anti-mouse IgG antibodies conjugated with different Alexa FluoR (Invitrogen) diluted 1:2000 in PBS. Sections were washed, mounted in Fluoromount and examined with a Leica SP8 confocal microscope. Detection of mitochondrial superoxide was performed in acute brain slices (400 μm) from 6 month-old mice by incubation with 5μM MitoSOX Red (Life Technologies) in ACSF for 10 min. Slices were then fixed with 4% paraformaldehyde at 4°C for 1 h. After washing and permeabilization in PBS, slices were further incubated with TO-PRO-3 Iodide (diluted in PBS, 1:2000, Life Technologies) at RT for 10 min, rinsed and mounted onto slides with Fluoromount. The slices were examined using a Leica SP8 confocal microscope with a 40X objective.

### Cell death and neurite length assays in response to Aβ

Aβ-induced neurotoxicity has been shown to correlate with the extent of beta sheet structure in Aβ oligomers (Simmons *et al*, 1994). These were formed by 24h incubation at 37°C of a 1mg/ml solution of Aβ 1-42 peptide (Sigma) in PBS. Aβ oligomers were added to cortical neuron cultures at different concentrations and the cultured were maintained at 37°C in a CO2 atmosphere. Cell death was assessed after overnight incubation by the appearance of cleaved caspase-3 positive neurons as detected by immunocytochemistry. Neurite length was assessed after 24h incubation by MAP2 immunostaining followed by image analysis using Image J Software.

### Internalization assay and super-resolution microscopy

Primary antibody incubation was performed using mouse 6E10 monoclonal antibody (Biolegend) against human Aβ diluted (1:200) in artificial cerebrospinal fluid (ACSF: 124mM NaCl, 3.7mM KCl, 1.0mM MgSO4, 2.5mM CaCl2, 1.2mM KH2PO4, 24.6mM NaHCO3, and 10mM D-glucose) for 1 hour at 4°C to label surface hAPP. For p75^NTR^ internalization, GT15057 antibody (Neuromics) against the receptor extracellular domain was used instead. The cultures were then washed in ACSF and incubated at 37°C to allow internalization. At different time points, the internalization was stopped by quick wash in 70% formic acid. Total staining (100%) was determined by direct fixation after antibody feeding, without acid wash. Baseline (t=0 min) was obtained by acid wash directly after antibody feeding without 37°C incubation. The cultures were then fixed by addition of 4% PFA in PBS for 15 min at room temperature, washed with PBS, and incubated in blocking buffer for 30 min at room temperature. Labelled hAPP and p75^NTR^ were detected by incubation with appropriate secondary antibodies at 1:2000 dilution in blocking buffer for 1h at room temperature. Cells were washed three times with PBS and mounted in Flouromount. In all experiments, cells were visualized on a Leica SP8 confocal microscope.

To assess co-localization of internalized APP with BACE1 by super-resolution microscopy, fixed coverslips were immunostained for BACE-1 prior to incubation with secondary antibodies (Alexa 555 for 6E10, Alexa 647 for BACE1, 1:2000). Imaging was performed on a Leica TCS SP8 X microscope. Super-resolution was done by the Leica HyVolution technique which can resolve down to 130nm. Images were captured using a 63x/1.4NA oil immersion objective and 4X zoom. Ten super-resolution images were acquired for each group. The colocalization analysis was done with Fiji software (NIH Image).

### Proximity ligation assay (PLA)

Hippocampal neurons were fixed for 15 min in 4% PFA, permeabilized, and blocked in 10% normal donkey serum and 0.3% Triton X-100 in PBS. Cells were then incubated overnight at 4°C with rabbit anti-p75^NTR^ (1:200, AB1554), mouse anti-human Aβ (6E10; 1:1000), and chicken anti-MAP2 (1:2000) antibodies in PBS supplemented with 3% BSA. The Duolink In Situ Proximity Ligation kit (Sigma) was used as per the manufacturer’s instructions with fluorophore-conjugated secondary antibody to recognize MAP2 (1:2000) included during the amplification step. The cultures were imaged with a Leica SP8 confocal microscope to detect PLA signals. PLA puncta were quantified using ImageJ software. Internalization of PLA signals followed the protocol described in the previous section, except that live neuron cultures were simultaneously fed with antibodies against APP (6E10) and p75^NTR^ (AB1554). MAP2 counterstaining was then performed after internalization was completed and following fixation of the cultures.

### Image analysis

For each mouse, five brain coronal sections spaced 120 μm were quantified. In each brain section, mosaic images in hippocampus were captured. ImageJ software was used to quantify positive signal area for the different markers. For Aβ, GFAP and Iba1, percentage of positive signal area was normalized to total area of hippocampus. For MitoSox staining, mean fluorescence intensity in a fixed area within the hippocampal pyramidal cell layer was quantified. For RTN3, positive signal area within Aβ plaques was for quantification and expressed as percentage of total Aβ plaque area. For quantification of MitoSox signal, stacks of 22 consecutive confocal images taken at 0.5μm intervals were acquired sequentially with two lasers (405 nm for TO-PRO and 561 nm for MitoSOX). All parameters were held constant for all the sections. Four images covering the pyramidal layer visualized with TO-PRO nuclear staining in CA1 hippocampal formation were captured for each mouse (three age-matched mice per genotype). Image J Software was used to quantify the mean fluorescence intensity in the pyramidal cell layer.

### Electrophysiology

Our electrophysiological procedures are described in greater detail in Shetty et al. (Shetty *et al*, 2015). Briefly, mice were decapitated after anesthesia with CO2 and the brains were quickly removed into cold (4°C) artificial cerebrospinal fluid (ACSF: 124mM NaCl, 3.7mM KCl, 1.0mM MgSO4, 2.5mM CaCl2, 1.2mM KH2PO4, 24.6mM NaHCO3, and 10mM D-glucose) equilibrated with 95% O2/5% CO2 (carbogen; total consumption 16L/h). From each mouse, 6-8 transverse hippocampal slices (400 μm-thick) were prepared from the right hippocampus by using a manual tissue chopper. Slices were incubated at 32°C in an interface chamber (Scientific System Design) at an ACSF flow rate of 1 mL/minute. One monopolar, lacquer coated, stainless steel electrode (5 MΩ; AM Systems) was positioned at an adequate distance within the stratum radiatum of the CA1 region for stimulating synaptic inputs of one neuronal population, thus evoking field excitatory post-synaptic potential (fEPSP) from Schaffer collateral-commissural-CA1 synapses. For recording, another electrode was placed in the CA1 apical dendritic layer. The signals were amplified by a differential amplifier (Model 1700, AM Systems) and were digitized using a CED 1401 analog-to-digital converter (Cambridge Electronic Design). After the pre-incubation period of 2 hours, an input-output curve (afferent stimulation vs fEPSP slope) was plotted prior to experiments. To set the test stimulus intensity, a fEPSP of 40% of its maximal amplitude was determined. Biphasic constant current pulses were used for stimulation. Late long-term potentiation (L-LTP) was induced using a theta burst stimulation (TBS) protocol which consists of 50 bursts (consisting of 4 stimuli) at an inter-stimulus interval of 10ms. The 50 bursts were applied over a period of 30s at 5 Hz (or at an inter-burst interval of 200ms). The slopes of fEPSPs were monitored online. The baseline was recorded for 30min. For baseline recording and testing at each time point, four 0.2 Hz biphasic constant-current pulses (0.1ms/polarity) were used. fEPSPs were recorded every 5min from 30min before stimulation up to 240min after stimulation across the CA1-CA3 Schaffer collaterals and normalized against t=0.

### Behavior tests

The novel object recognition (NOR) test for mice consists of 3 days of exposure training, followed by a short term memory (STM) test 20min after training, a long term memory (LTM) test 24h later and a remote memory test 2 weeks later. The objects are chosen based on similarities in dimensions and complexity. Tests are carried out in an acrylic box (20.32×40.5×16 cm L×W×H) that is sanitized, together with the objects, with 70% ethanol between each experiment. The time spent with an object includes direct visual orientation towards an object within half a body length of the object, sniffing, touching or climbing on the object. The tests are video recorded and preference scores are calculated as time spent with novel object minus time spent with familiar object divided by the total time spent with both objects. Positive scores indicate preferences for the novel object; negative scores show preferences for the familiar object.

For the Barnes maze spatial memory test, spatial cues were placed around the maze and these were kept constant throughout the study. On the first day of training, the mouse was placed in the escape box for one minute. The animal was then placed in the center of the maze inside a black chamber. As in all subsequent sessions, the chamber was removed after 10 seconds, whereupon a buzzer (80 dB) and a light (400 lux) were turned on, and the mouse was free to explore the maze for 3 min or until the mouse entered the escape tunnel. The tunnel was always located underneath the same hole, which was randomly determined for each mouse. The platform was moved every day by 90° to avoid any odorant cue but the spatial cues and the tunnel position remained the same. Mice were trained using this protocol once daily for 4 days. For the test sessions, the escape tunnel was removed, and the mouse was allowed to freely explore the maze for a maximum of 3 min to assess spatial memory. The short-term memory test was conducted 20-30 min after the first training. Long-term and remote memory tests were conducted 24h and 14 days after the last training, respectively. To quantify the preference for the trained target quadrant, time spent in the target quadrant and time spent in the other quadrants was measured.

### Statistical analysis

Statistics analyses were performed using Prism 7 software (GraphPad, SPSS IBM corporation) and Microsoft Excel (Microsoft). Results are presented as mean ± standard error of the mean (SEM). Student’s t test, one-way ANOVA or two-way ANOVA were performed to test statistical significance according the requirements of the experiment. One-way ANOVA followed by post hoc test was used for statistical analysis of immunohistopathology data. Behavioral data were analyzed using two-way ANOVA with post hoc analysis. Statistical significance: *, p<0.05; **, p<0.01 and ***, p<0.001.

## Acknowledgements

We thank Peiyan Wong for assistance with behavioral studies. This research was funded by grants NMRC/CBRG/0107/2016 (to C.F.I.) and NMRC/OFIRG/0037/2017 (to S.S.) from the Singapore National Medical Research Council, and MOE2018-T2-1-129 (to C.F.I.) and MOE2017-T3-1-002 (to S.S.), from Singapore Ministry of Education, ODPRT Strategic Programme Award (to C.F.I. and S.S.) and Aspiration Fund World Class (to C.F.I.) from the National University of Singapore, and VR-2016-01538 (to C.F.I.) from the Swedish Research Council (to C.F.I.).

## Author contributions

C.Y. and K.Y.G. performed all experiments, except cell death assay, performed by K.T., and electrophysiology, performed by L.W.W. C.Y. and C.F.I. designed the experiments. C.F.I. and S.S. directed research. C.F.I. wrote the manuscript.

## Supplementary information

Supplementary Figures 1 to 6 Table S1

## References

Bronfman FC, Tcherpakov M, Jovin TM & Fainzilber M (2003) Ligand-induced internalization of the p75 neurotrophin receptor: a slow route to the signaling endosome. J Neurosci 23: 3209–3220

Buxbaum JD, Thinakaran G, Koliatsos V, O’Callahan J, Slunt HH, Price DL & Sisodia SS (1998) Alzheimer Amyloid Protein Precursor in the Rat Hippocampus: Transport and Processing through the Perforant Path. Journal of Neuroscience 18: 9629–9637

Carey RM, Balcz BA, Lopez-Coviella I & Slack BE (2005) Inhibition of dynamin-dependent endocytosis increases shedding of the amyloid precursor protein ectodomain and reduces generation of amyloid beta protein. BMC Cell Biol. 6: 30–10

Chakravarthy B, Gaudet C, Ménard M, Atkinson T, Brown L, Laferla FM, Armato U & Whitfield J (2010) Amyloid-beta peptides stimulate the expression of the p75(NTR) neurotrophin receptor in SHSY5Y human neuroblastoma cells and AD transgenic mice. J. Alzheimers Dis. 19: 915–925

Chakravarthy B, Ménard M, Ito S, Gaudet C, Dal-Pra I, Armato U & Whitfield J (2012) Hippocampal membrane-associated p75NTR levels are increased in Alzheimer’s disease. J. Alzheimers Dis. 30: 675–684

Das U, Scott DA, Ganguly A, Koo EH, Tang Y & Roy S (2013) Activity-induced convergence of APP and BACE-1 in acidic microdomains via an endocytosis-dependent pathway. Neuron 79: 447–460

Das U, Wang L, Ganguly A, Saikia JM, Wagner SL, Koo EH & Roy S (2016) Visualizing APP and BACE-1 approximation in neurons yields insight into the amyloidogenic pathway. Nat Neurosci 19: 55–64

Dikalov SI & Harrison DG (2014) Methods for detection of mitochondrial and cellular reactive oxygen species. Antioxid. Redox Signal. 20: 372–382

Ernfors P, Lindefors N, Chan-Palay V & Persson H (1990) Cholinergic neurons of the nucleus basalis express elevated levels of nerve growth factor receptor mRNA in senile dementia of the Alzheimer type. Dementia 1: 138–145

Fombonne J, Rabizadeh S, Banwait S, Mehlen P & Bredesen DE (2009) Selective vulnerability in Alzheimer’s disease: amyloid precursor protein and p75(NTR) interaction. Ann. Neurol. 65: 294–303

Goh ETH, Lin Z, Ahn BY, Lopes-Rodrigues V, Dang N-H, Salim S, Berger B, Dymock B, Senger DL & Ibáñez CF (2018) A Small Molecule Targeting the Transmembrane Domain of Death Receptor p75NTRInduces Melanoma Cell Death and Reduces Tumor Growth. Cell Chemical Biology 25: 1485–1494.e5 Available at: http://www.cell.com.libproxy1.nus.edu.sg/article/S2451945618303027/fulltext

Haass C, Kaether C, Thinakaran G & Sisodia S (2012) Trafficking and Proteolytic Processing of APP. Cold Spring Harbor Perspectives in Medicine 2: a006270–a006270

Hu X, Shi Q, Zhou X, He W, Yi H, Yin X, Gearing M, Levey A & Yan R (2007) Transgenic mice overexpressing reticulon 3 develop neuritic abnormalities. EMBO J 26: 2755–2767

Hu X-Y, Zhang H-Y, Qin S, Xu H, Swaab DF & Zhou J-N (2002) Increased p75(NTR) expression in hippocampal neurons containing hyperphosphorylated tau in Alzheimer patients. Exp Neurol 178: 104–111

Ibáñez CF & Simi A (2012) p75 neurotrophin receptor signaling in nervous system injury and degeneration: paradox and opportunity. Trends Neurosci 35: 431–440

Karran E, Mercken M & De Strooper B (2011) The amyloid cascade hypothesis for Alzheimer’s disease: an appraisal for the development of therapeutics. Nat Rev Drug Discov 10: 698–712

Knowles JK, Rajadas J, Nguyen T-VV, Yang T, LeMieux MC, Griend LV, Ishikawa C, Massa SM, Wyss-Coray T & Longo FM (2009) The p75 Neurotrophin Receptor Promotes Amyloid-beta(1-42)-Induced Neuritic Dystrophy In Vitro and In Vivo. J Neurosci 29: 10627–10637

Koch G, Di Lorenzo F, Bonnì S, Ponzo V, Caltagirone C & Martorana A (2012) Impaired LTP-but not LTD-Like Cortical Plasticity in Alzheimer’s Disease Patients. Journal of Alzheimer’s Disease 31: 593–599

Koo EH & Squazzo SL (1994) Evidence that production and release of amyloid beta-protein involves the endocytic pathway. J Biol Chem 269: 17386–17389

Kwart D, Gregg A, Scheckel C, Murphy E, Paquet D, Duffield M, Fak J, Olsen O, Darnell R & Tessier-Lavigne M (2019) A Large Panel of Isogenic APP and PSEN1 Mutant Human iPSC Neurons Reveals Shared Endosomal Abnormalities Mediated by APP β-CTFs, Not Aβ. Neuron 0: 256–270.e5

Lauritzen I, Pardossi-Piquard R, Bauer C, Brigham E, Abraham J-D, Ranaldi S, Fraser P, St-George-Hyslop P, Le Thuc O, Espin V, Chami L, Dunys J & Checler F (2012) The β-secretase-derived C-terminal fragment of βAPP, C99, but not Aβ, is a key contributor to early intraneuronal lesions in triple-transgenic mouse hippocampus. J Neurosci 32: 16243–1655a

Lauritzen I, Pardossi-Piquard R, Bourgeois A, Pagnotta S, Biferi M-G, Barkats M, Lacor P, Klein W, Bauer C & Checler F (2016) Intraneuronal aggregation of the β-CTF fragment of APP (C99) induces Aβ-independent lysosomal-autophagic pathology. Acta Neuropathol. 132: 257–276

Lee KF, Li E, Huber LJ, Landis SC, Sharpe A, Chao MV & Jaenisch R (1992) Targeted Mutation of the Gene Encoding the Low Affinity Ngf Receptor P75 Leads to Deficits in the Peripheral Sensory Nervous-System. Cell 69: 737–749

Liepinsh E, Ilag LL, Otting G & Ibáñez CF (1997) NMR structure of the death domain of the p75 neurotrophin receptor. EMBO J 16: 4999–5005

Lu B, Nagappan G, Guan X, Nathan PJ & Wren P (2013) BDNF-based synaptic repair as a disease-modifying strategy for neurodegenerative diseases. Nature Reviews Neuroscience 14: 401–416

Mango D, Saidi A, Cisale GY, Feligioni M, Corbo M & Nisticò R (2019) Targeting Synaptic Plasticity in Experimental Models of Alzheimer’s Disease. Front Pharmacol 10: 778

Mufson EJ & Kordower JH (1992) Cortical neurons express nerve growth factor receptors in advanced age and Alzheimer disease. Proc Natl Acad Sci USA 89: 569–573

Oakley H, Cole SL, Logan S, Maus E, Shao P, Craft J, Guillozet-Bongaarts A, Ohno M, Disterhoft J, Van Eldik L, Berry R & Vassar R (2006) Intraneuronal beta-amyloid aggregates, neurodegeneration, and neuron loss in transgenic mice with five familial Alzheimer’s disease mutations: potential factors in amyloid plaque formation. J Neurosci 26: 10129–10140

Palop JJ & Mucke L (2010) Amyloid-beta-induced neuronal dysfunction in Alzheimer’s disease: from synapses toward neural networks. Nat Neurosci 13: 812–818

Pera M, Larrea D, Guardia-Laguarta C, Montesinos J, Velasco KR, Agrawal RR, Xu Y, Chan RB, Di Paolo G, Mehler MF, Perumal GS, Macaluso FP, Freyberg ZZ, Acin-Perez R, Enriquez JA, Schon EA & Area-Gomez E (2017) Increased localization of APP-C99 in mitochondria-associated ER membranes causes mitochondrial dysfunction in Alzheimer disease. EMBO J 36: 3356–3371

Perini G, Della-Bianca V, Politi V, Valle Della G, Dal-Pra I, Rossi F & Armato U (2002) Role of p75 neurotrophin receptor in the neurotoxicity by beta-amyloid peptides and synergistic effect of inflammatory cytokines. J Exp Med 195: 907–918

Rabizadeh S, Bitler CM, Butcher LL & Bredesen DE (1994) Expression of the low-affinity nerve growth factor receptor enhances beta-amyloid peptide toxicity. Proc Natl Acad Sci U S A 91: 10703–10706

Selkoe DJ (2002) Alzheimer’s disease is a synaptic failure. Science 298: 789–791

Selkoe DJ & Hardy J (2016) The amyloid hypothesis of Alzheimer’s disease at 25 years. EMBO Mol Med 8: 595–608

Selkoe DJ, Yamazaki T, Citron M, Podlisny MB, Koo EH, Teplow DB & Haass C (1996) The role of APP processing and trafficking pathways in the formation of amyloid beta-protein. Ann N Y Acad Sci 777: 57–64

Sherman MA & Lesné SE (2011) Detecting aβ*56 oligomers in brain tissues. Methods Mol. Biol. 670: 45–56

Shetty MS, Sharma M, Hui NS, Dasgupta A, Gopinadhan S & Sajikumar S (2015) Investigation of Synaptic Tagging/Capture and Cross-capture using Acute Hippocampal Slices from Rodents. J Vis Exp: e53008–e53008

Simmons LK, May PC, Tomaselli KJ, Rydel RE, Fuson KS, Brigham EF, Wright S, Lieberburg I, Becker GW & Brems DN (1994) Secondary structure of amyloid beta peptide correlates with neurotoxic activity in vitro. Mol Pharmacol 45: 373–379

Sotthibundhu A, Sykes AM, Fox B, Underwood CK, Thangnipon W & Coulson EJ (2008) beta-amyloid(1-42) induces neuronal death through the p75 neurotrophin receptor. J Neurosci 28: 3941–3946

Swerdlow RH (2018) Mitochondria and Mitochondrial Cascades in Alzheimer’s Disease. Journal of Alzheimer’s Disease 62: 1403–1416

Tanaka K, Kelly CE, Goh KY, Lim KB & Ibáñez CF (2016) Death Domain Signaling by Disulfide-Linked Dimers of the p75 Neurotrophin Receptor Mediates Neuronal Death in the CNS. J Neurosci 36: 5587–5595

Underwood CK & Coulson EJ (2008) The p75 neurotrophin receptor. Int J Biochem Cell Biol 40: 1664–1668

Vassar R, Kovacs DM, Yan R & Wong PC (2009) The beta-secretase enzyme BACE in health and Alzheimer’s disease: regulation, cell biology, function, and therapeutic potential. J Neurosci 29: 12787–12794

Vilar M, Charalampopoulos I, Kenchappa RS, Simi A, Karaca E, Reversi A, Choi S, Bothwell M, Mingarro I, Friedman WJ, Schiavo G, Bastiaens PIH, Verveer PJ, Carter BD & Ibáñez CF (2009) Activation of the p75 neurotrophin receptor through conformational rearrangement of disulphide-linked receptor dimers. Neuron 62: 72–83

Wang Y-J, Wang X, Lu J-J, Li Q-X, Gao C-Y, Liu X-H, Sun Y, Yang M, Lim Y, Evin G, Zhong J-H, Masters C & Zhou X-F (2011) p75NTR regulates Abeta deposition by increasing Abeta production but inhibiting Abeta aggregation with its extracellular domain. J Neurosci 31: 2292–2304

Xu W, Weissmiller AM, White JA, Fang F, Wang X, Wu Y, Pearn ML, Zhao X, Sawa M, Chen S, Gunawardena S, Ding J, Mobley WC & Wu C (2016) Amyloid precursor protein-mediated endocytic pathway disruption induces axonal dysfunction and neurodegeneration. J Clin Invest 126: 1815–1833

Yaar M, Zhai S, Pilch PF, Doyle SM, Eisenhauer PB, Fine RE & Gilchrest BA (1997) Binding of beta-amyloid to the p75 neurotrophin receptor induces apoptosis. A possible mechanism for Alzheimer’s disease. J Clin Invest 100: 2333–2340

